# Fission yeast cells use distinct cell size control mechanisms for size adaptation to osmotic, oxidative, or low glucose conditions

**DOI:** 10.1101/2025.11.04.686600

**Authors:** Elena J. Cabral, Pablo Andres, Geraldin Argandona, Paige Duggan, Benjamin M. Kuran, Kristi E. Miller

## Abstract

Cells maintain an appropriate size to function, yet the mechanisms that enable size adaptation to environmental stress remain poorly understood. Fission yeast cells enter mitosis and divide at a threshold size when cyclin-dependent kinase (Cdk1) is activated through size- and time-dependent scaling of its regulators: Cdr2 kinase with cell surface area, Cdc25 phosphatase with cell volume, and mitotic cyclin Cdc13 with cell cycle time. This integrated size control network is characterized in nutrient-rich conditions, but under stress it remains unclear which size parameters cells monitor, and which size- or time-sensing pathways mediate adaptation. Using high-throughput image analysis, we quantified the geometry of dividing cells under osmotic, oxidative, and low glucose conditions. Wild-type cells increased their surface area-to-volume (SA:Vol) ratio in low glucose but decreased it under osmotic or oxidative stress, revealing distinct geometric strategies for environmental size adaptation. Genetic perturbations of size- and time-sensing pathways revealed that Cdc25 is required for volume-based adaptation to oxidative and osmotic stress, Cdc13 contributes to osmotic stress response, and Cdr2 promotes surface area-based expansion in low glucose. Although disrupting individual pathways altered normal geometric responses, cells remained viable, suggesting that a modular size control system enables flexible geometric adaptation to changing environments.

## INTRODUCTION

Cell size is a fundamental property that influences metabolism and gene expression and must be precisely regulated to ensure proper cell function. Most cells control their size by delaying cell cycle transitions until they reach a threshold size, indicating that growth and cell cycle progression are tightly coordinated (Turner *et al*., 2012: Amodeo and Skotheim, 2016). However, cell size is not fixed. Environmental factors influence the size of cells in both unicellular and multicellular organisms. For example, starvation leads to reduced cell size in *Drosophila*, rats, and yeast (Robertson, 1963; Hirsch and Han, 1969; Fantes and Nurse, 1977; Johnston *et al*., 1979). The ability to adjust cell size in response to environmental conditions is critical to maintain cellular function and survival, yet the molecular mechanisms that enable cells to adapt their size remain poorly understood (Kellogg and Levin, 2022).

Fission yeast *Schizosaccharomyces pombe* is a powerful model system to investigate size control mechanisms due to its genetic tractability and simple rod shape. Fission yeast cells grow by linear extension and divide at a reproducible size that has historically been measured by cell length but is now more accurately defined by surface area (SA) (Mitchison and Nurse, 1985; Pan *et al*., 2014; Facchetti *et al*., 2019). The core regulators of mitotic entry and cell size in fission yeast are conserved across eukaryotes (Figure 1A). Entry into mitosis is triggered by activation of the cyclin-dependent kinase Cdk1 (also called Cdc2 in fission yeast) bound to its B-type cyclin, Cdc13. Cdk1 activity is inhibited by the kinase Wee1 in small cells and activated by the phosphatase Cdc25 once cells reach a threshold size (Russell and Nurse, 1986; Gould and Nurse, 1989). Recent work has shown that Cdk1 is activated through size- and time-dependent scaling of its regulators: the kinase Cdr2, which inhibits Wee1, scales with cell surface area (Pan *et al*., 2014; Facchetti *et al*., 2019); the phosphatase Cdc25 scales with cell volume; and the cyclin Cdc13 accumulates over cell cycle time (Miller *et al*., 2023) (Figure 1B). These size- and time-dependent properties of Cdk1 activators define a modular size control network that coordinates mitotic entry with cell growth through multiple pathways.

**Figure 1:**
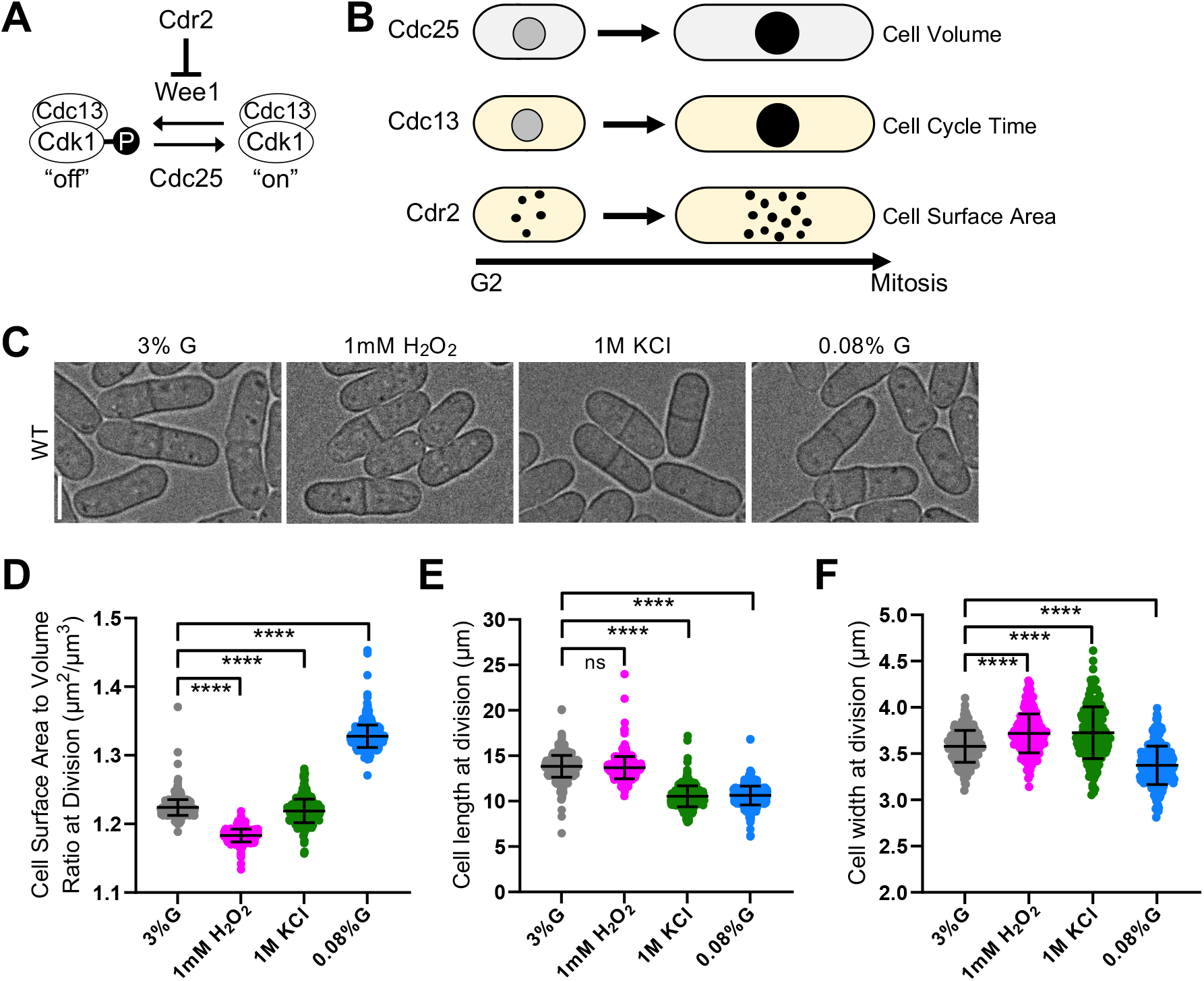
WT cells adjust their SA:Vol ratio based on stress condition. (A) Schematic of biochemical pathway regulating mitotic entry and cell size. (B) Cdk1 activators scale with distinct aspects of size or time. (C) Brightfield images of WT cells grown under various environmental conditions. (D) SA:Vol ratio at division. \*\*\*\**p* < 0.0001. *n* > 300 for each condition. (E) Cell length at division. ns, not significant; \*\*\*\**p* < 0.0001. *n* > 300 for each condition. (F) Cell width at division. \*\*\*\**p* < 0.0001. *n* > 200 for each condition.

While these cell surface area-(Cdr2), volume-(Cdc25), and time-sensing (Cdc13) pathways have been characterized under nutrient-rich conditions in fission yeast (Pan *et al*., 2014; Facchetti *et al*., 2019; Miller *et al*., 2023), their roles under environmental stress remain unclear. Environmental conditions such as glucose limitation and osmotic or oxidative stress profoundly alter growth, morphology, and cell size (Kellogg and Levin, 2022; Chadha *et al*., 2024), yet it remains unknown which geometric parameter (surface area, volume, or others) cells monitor to divide or which size- or time-sensing pathways drive adaptation. Understanding how cells adjust or reweight these pathways during stress will reveal how size control is maintained to preserve cell function in changing environments.

Here, we investigate how fission yeast cells adapt their geometry and size control pathways under three distinct stress conditions: low glucose, oxidative stress, and osmotic stress. Using a high-throughput image analysis pipeline (Miller *et al*., 2023), we quantified cell geometry in wild-type (WT) and mutant strains in which surface area-, volume-, or time-sensing pathways were disrupted. We show that WT cells adopt distinct geometric strategies: increasing their surface area-to-volume (SA:Vol) ratio under low glucose but decreasing the SA:Vol ratio under oxidative or osmotic stress. Genetic perturbations of *cdc25, cdc13*, and *cdr2* reveal that each pathway makes unique, stress-specific contributions to geometric adaptation. Our findings support a model in which fission yeast flexibly engage different pathways to adjust cell geometry in response to environmental stress. Although disrupting individual size- and time-sensing pathways alters normal geometric adaptation, cells remain viable, highlighting the robustness and modularity of the size control network. This modularity likely provides an adaptive advantage, allowing cells to sustain division and maintain homeostasis under changing environmental conditions.

## RESULTS AND DISCUSSION

To investigate how distinct cell size control pathways contribute to size adaptation under environmental stress, we analyzed fission yeast mutants in which the surface area-, volume-, or time-sensing pathways were disrupted (described below), along with a WT control. Cells were grown for 48 hours in YE4S medium containing 3% glucose as a control or under stress-inducing conditions: low glucose (0.08% glucose), oxidative stress (1mM H_2_O_2_), or osmotic stress (1M KCl) to capture long-term rather than transient stress responses. All strains expressed a BFP-NLS nuclear marker to identify dividing cells, and brightfield images were collected to visualize cell boundaries for cell size measurements. Cell size was quantified using an image analysis pipeline, in which binary masks generated in ImageJ were analyzed with MATLAB codes to obtain nuclear and cell dimensions (Miller *et al*., 2023). From these data, we obtained cell width, length, surface area, and volume of dividing cells, and calculated the SA:Vol ratio. For each genotype, cell geometric parameters were compared across stress conditions relative to the 3% glucose control using one-way ANOVA. These analyses provided a quantitative framework to determine how the Cdc25 (volume), Cdc13 (time), and Cdr2 (surface area) pathways contribute to stress-dependent changes in cell geometry.

### Wild-type cells exhibit stress-specific adaptations in SA:Vol ratio

First, we examined how WT cells alter their geometry in response to environmental stress (Figure 1C). Under oxidative and osmotic stress, the SA:Vol ratio decreased, indicating greater cell volume expansion relative to surface area (Figure 1D). These changes occurred through stress-specific alterations in cell dimensions. During oxidative stress, overall cell size remained relatively constant, with cells becoming wider but maintaining their length (Figures 1E and 1F). Under osmotic stress, overall cell size decreased through reduced length accompanied by a modest increase in width (Figures 1E and 1F). In contrast, under low glucose conditions, cells increased their SA:Vol ratio, indicating a shift toward maximizing surface area relative to volume (Figure 1D). WT cells reduced both their length and width in low glucose (Figures 1E and 1F). While previous studies primarily examined cell length changes under stress, our stress-specific geometric adaptations are consistent with prior observations, including no major cell size changes under oxidative stress (Degols, *et al*., 1996; Salat-Canela *et al*., 2021), reduced cell length under osmotic stress (Shiozaki and Russell, 1995), and coordinated changes in length, width, and SA:Vol ratio under low glucose conditions (Fantes and Nurse, 1977; Bertaux *et al*., 2023).

These results show that WT cells adopt environment-specific geometric changes, preferably expanding in volume under oxidative and osmotic stress and in surface area under glucose limitation. Subtle adjustments in the SA:Vol ratio may reflect physiological tuning that optimizes nutrient uptake, waste removal, and environmental sensing that supports adaptation to different environments (Amodeo and Skotheim, 2016).

### Cdc25 is required for cell volume expansion under oxidative and osmotic stress

To determine how the Cdc25 volume-sensing pathway contributes to size adaptation under stress, we examined mutants that disrupt either Cdc25 protein abundance or its size-dependent expression. Cdc25 is a mitotic activator whose nuclear concentration scales with cell volume, thereby linking cell volume to mitotic entry (Miller *et al*., 2023). We hypothesized that disrupting this scaling relationship might impair stress-dependent volume expansion observed under oxidative and osmotic stress.

To reduce Cdc25 protein levels, we used a degron-DaMP tagged allele of *cdc25* (Breslow *et al*., 2008). Under oxidative and osmotic stress, *cdc25-degron-DaMP* cells failed to maximize volume as observed in WT cells (Figure 2A and 2B). Instead, in oxidative stress these mutant cells maintained a SA:Vol ratio similar to the 3% glucose control and increased in size primarily by elongating, while cell width remained unchanged (Figures 2B-D). Under osmotic stress, *cdc25-degron-DaMP* cells increased their SA:Vol ratio, opposite to the decrease observed in WT cells, by elongating and decreasing in width (Figures 2B-D). In contrast, under low glucose conditions, *cdc25-degron-DaMP* cells adapted similar to WT, reducing length and increasing SA:Vol ratio (Figures 2B-D).

**Figure 2:**
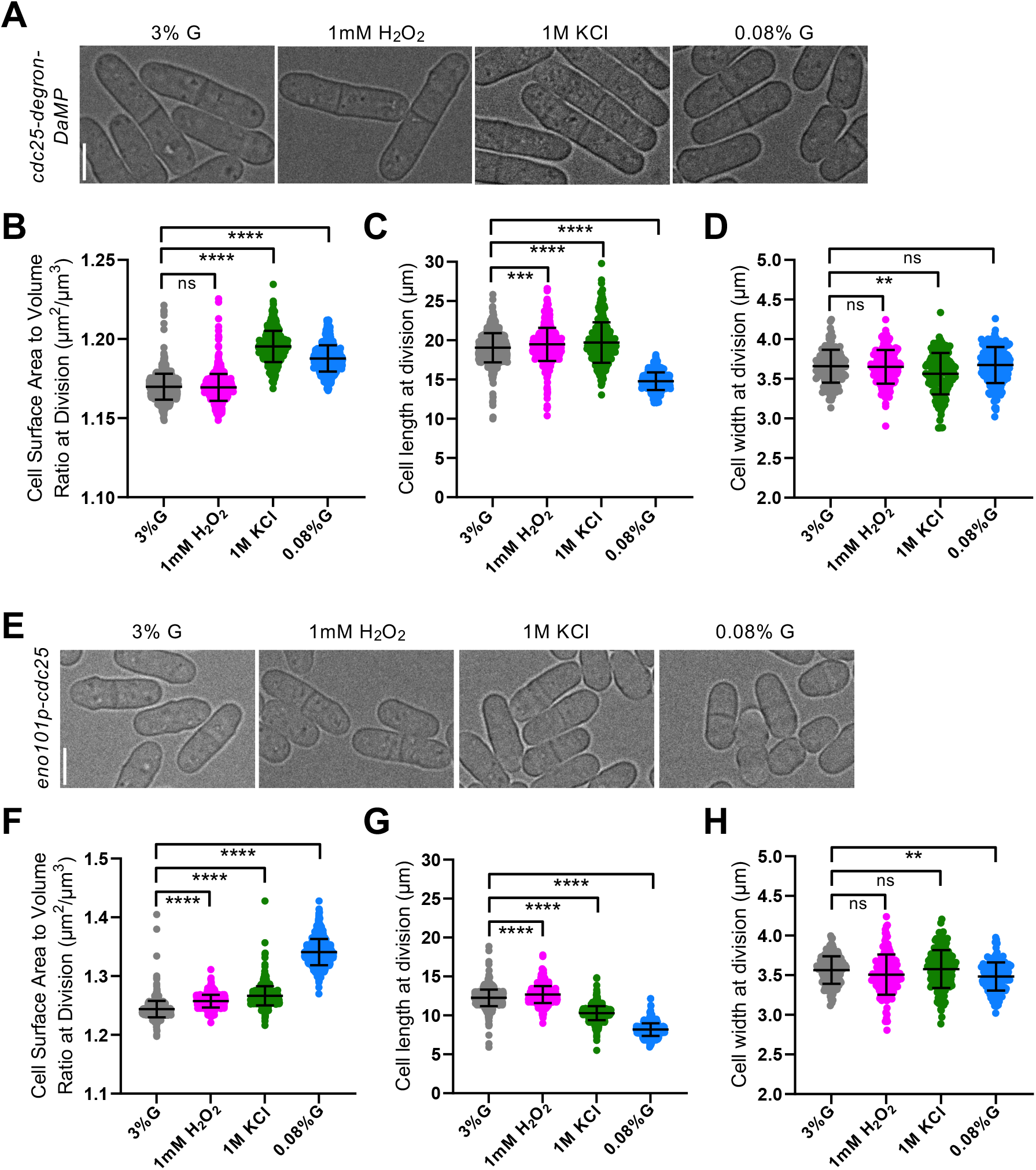
Cdc25 is necessary for maximizing cell volume under oxidative and osmotic stress. (A) Brightfield images of *cdc25-degron-DaMP* cells grown under various environmental conditions. (B) SA:Vol ratio at division. ns, not significant; \*\*\*\**p* < 0.0001. *n* > 250 for each condition. (C) Cell length at division. \*\*\**p* < 0.001; \*\*\*\**p* < 0.0001. *n* > 250 for each condition. (D) Cell width at division. ns, not significant; \*\**p* < 0.01. *n* > 150 for each condition. (E) Brightfield images of *eno101p-cdc25* cells grown under various environmental conditions. (F) SA:Vol ratio at division. \*\*\*\**p* < 0.0001. *n* > 650 for each condition. (G) Cell length at division. \*\*\*\**p* < 0.0001. *n* > 650 for each condition. (H) Cell width at division. ns, not significant; \*\**p* < 0.01. *n* > 150 for each condition.

Because *cdc25-degron-DaMP* cells divide at a larger size than WT (Figure S1A), we next asked whether their altered geometry under oxidative and osmotic stress arose from disrupting Cdc25 volume-scaling or simply differences in division size. To distinguish between these possibilities, we analyzed the *eno101p-cdc25* strain, in which *cdc25* expression is driven by a constitutive *eno101* promoter (Wang *et al*., 2014) rather than its native, size-dependent promoter (Keifenheim *et al*., 2017). *eno101p-cdc25* cells exhibit strongly diminished size-dependent nuclear accumulation of Cdc25 (Figure S1B-D) and divide at a reduced cell size than WT (Figure S1A). Under oxidative and osmotic stress, *eno101p-cdc25* cells also failed to maximize volume and instead relied on surface area expansion (Figure 2E and 2F), similar to *cdc25-degron-DaMP* cells. During oxidative stress, cells elongated without changing width, while osmotic stress caused them to become shorter while maintaining width (Figures 2G and 2H). Under low glucose conditions, *eno101p-cdc25* cells mirrored the WT response, increasing their SA:Vol ratio and decreasing both length and width (Figure 2E-H).

Together, these results show that disrupting either Cdc25 protein abundance or size-dependent expression shifts cells from a volume-to a surface area-based adaptation strategy. Proper Cdc25 scaling with volume thus ensures coupling between cell volume and mitotic entry, allowing cells to adjust their size appropriately in response to oxidative or osmotic stress.

### Time sensing pathway (Cdc13) supports proper size adaptation under osmotic stress

We next investigated how perturbing the Cdc13 time-sensing pathway affects cell geometry under environmental stress. Mitotic cyclin Cdc13 nuclear accumulation scales with cell cycle duration, coupling time and mitotic entry (Miller *et al*., 2023). To test whether increasing *cdc13* dosage influences geometric adaptation under stress, we examined a *cdc13-2x* strain carrying an additional copy of *cdc13* integrated at an exogenous locus. *cdc13-2x* cells divided at a smaller size than WT, reflecting accelerated mitotic entry (Figure S1A).

Under oxidative stress, *cdc13-2x* cells adapted similarly to WT. These cells increased in both length and width, resulting in a decrease in the SA:Vol ratio (Figures 3A-D). Under low glucose conditions, *cdc13-2x* cells again responded like WT cells, decreasing length and showing a moderate increase in SA:Vol ratio (Figures 3B-D). In contrast, under osmotic stress, *cdc13-2x* cells failed to maximize cell volume (Figure 3B). These cells decreased in length and moderately increased in width, leading to an overall increase in SA:Vol ratio (Figure 3B-D), opposite to WT cells. This impaired volume expansion suggests that proper temporal control of mitotic entry through Cdc13 is required for size adaptation during osmotic stress. Although both oxidative and osmotic stress slow growth, cells maintain nearly constant size over time under oxidative stress, whereas osmotic stress causes a sustained reduction in size relative to time in G_2_ (Shiozaki and Russell, 1995; Degols et al., 1996; Salat-Canela *et al*., 2021), potentially increasing reliance on Cdc13’s time-sensing function to coordinate mitotic timing with growth. Together, the time-sensing (Cdc13) and volume-sensing (Cdc25) pathways likely maintain size homeostasis under osmotic stress.

**Figure 3:**
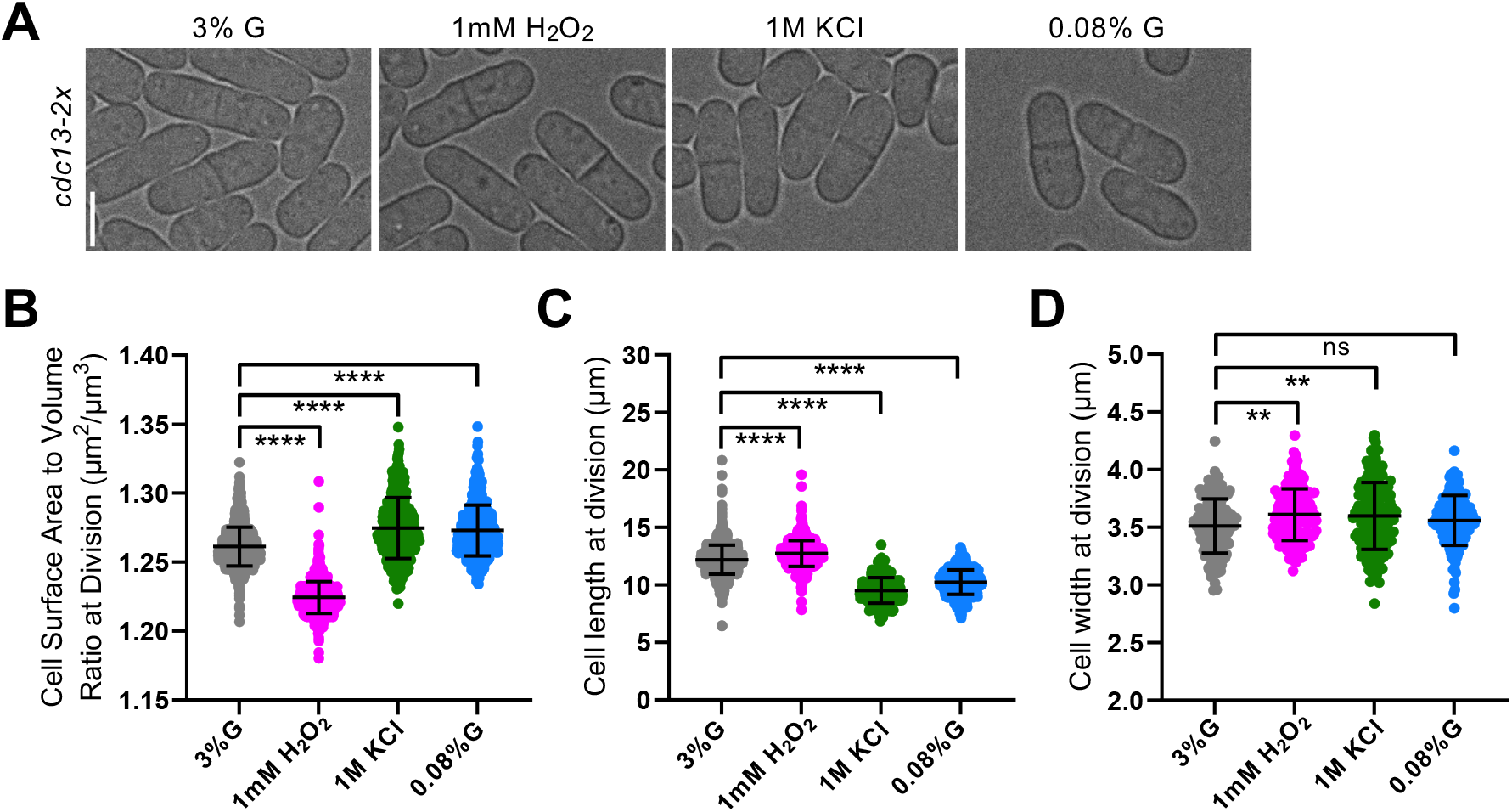
Cdc13 time-sensing pathway is necessary for proper size adaptation under osmotic stress. (A) Brightfield images of *cdc13-2x* cells grown under various environmental conditions. (B) SA:Vol ratio at division. \*\*\*\**p* < 0.0001. *n* > 425 for each condition. (C) Cell length at division. \*\*\*\**p* < 0.0001. *n* > 425 for each condition. (D) Cell width at division. ns, not significant; \*\**p* < 0.01. *n* > 150 for each condition.

### The Cdr2 cell surface area-sensing pathway supports surface area maximization under low glucose conditions

We next examined how perturbing the Cdr2 surface area-sensing pathway affects cell geometry adaptation under environmental stress. Cdr2 functions as a cortical scaffold that regulates Wee1 activity in relation to cell size (Allard *et al*., 2018) and enforces a surface area threshold for mitotic entry (Pan *et al*., 2014; Facchetti *et al*., 2019). We hypothesized that Cdr2 might be necessary for the surface area-based expansion observed under low glucose. To test this, we analyzed *cdr2Δ* cells (Figure 4A).

**Figure 4:**
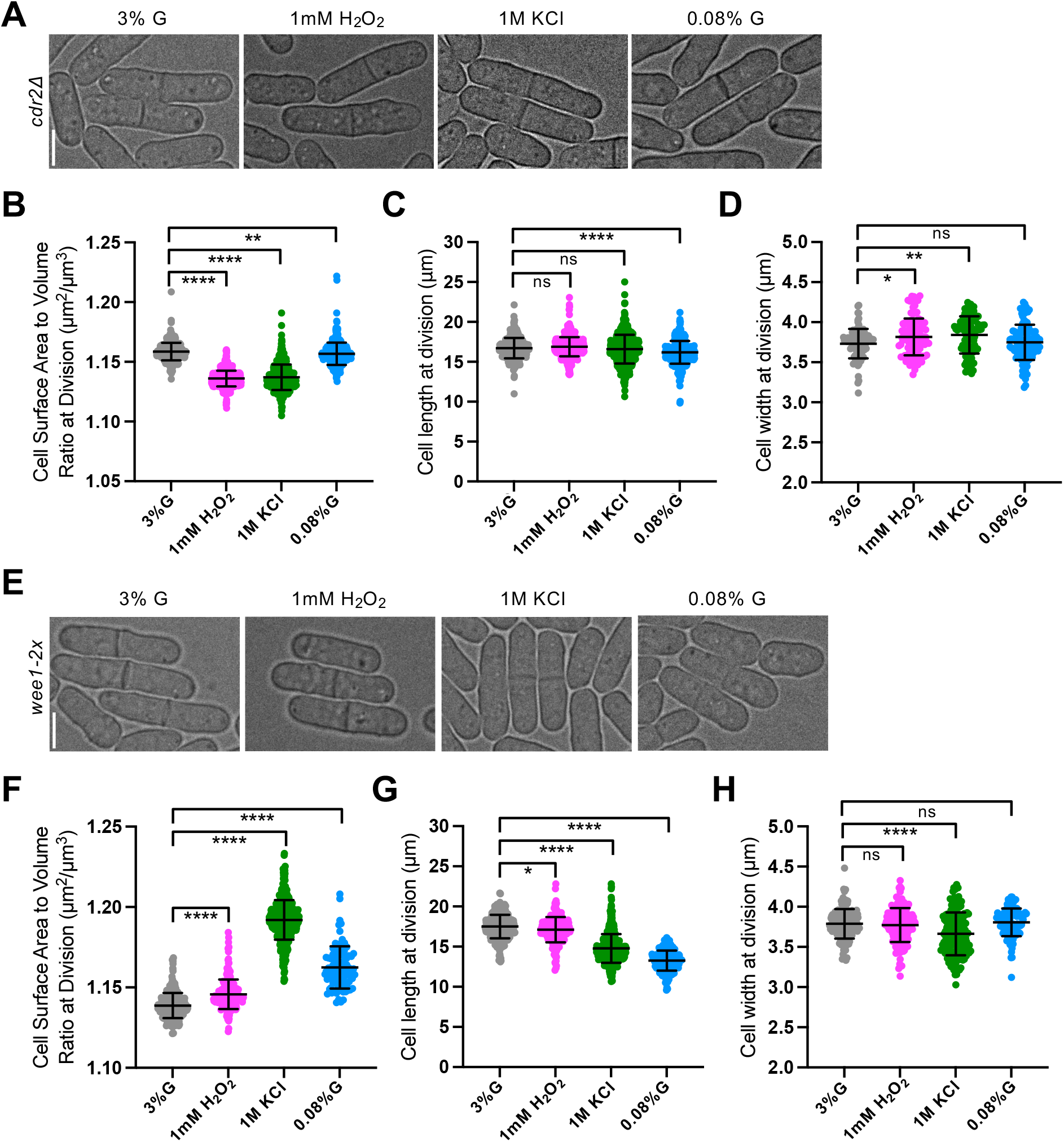
*cdr2Δ* cells fail to maximize surface area in low glucose, while *wee1-2x* mutants increase SA:Vol ratio under all stress conditions. (A) Brightfield images of *cdr2Δ* cells grown under various environmental conditions. (B) SA:Vol ratio at division \*\**p* < 0.01; \*\*\*\**p* < 0.0001. *n* > 400 for each condition. (C) Cell length at division. ns, not significant; \*\*\*\**p* < 0.0001. *n* > 400 for each condition. D) Cell width at division. ns, not significant; \**p* < 0.04; \*\**p* < 0.003; *n* > 100 for each condition. (E) Brightfield images of *wee1-2x* cells grown under various environmental conditions. (F) SA:Vol ratio at division. \*\*\*\**p* < 0.0001. *n* > 400 for each condition. (G) Cell length at division. \**p* < 0.03; \*\*\*\**p* < 0.0001. *n* > 400 for each condition. (H) Cell width at division. ns, not significant; \*\*\*\**p* < 0.0001. *n* > 100 for each condition.

Under oxidative and osmotic stress, *cdr2Δ* cells adapted similarly to WT cells, showing a decrease in SA:Vol ratio (Figure 4B). In both conditions, *cdr2Δ* cells displayed little to no change in cell length and a modest increase in width (Figure 4C and 4D), mirroring WT under oxidative stress (Figure 1D and 1E). These findings indicate that Cdr2 is dispensable for volume-based adaptation during oxidative and osmotic stress. In contrast, under low glucose conditions, *cdr2Δ* cells failed to maximize surface area (Figure 4B). Although they decreased in length slightly (Figure 4C), this decrease was smaller than observed in WT cells (0.52µm for *cdr2Δ* vs 3.2µm for WT), and cell width remained unchanged (Figure 4D). Consequently, *cdr2Δ* cells exhibited a reduction in SA:Vol ratio (Figure 4B), opposite to the increase observed in WT cells. These results suggest that Cdr2 promotes surface area-based expansion during glucose limitation, likely by enforcing a surface area threshold for division in low glucose environments.

To determine whether these effects reflect a specific role of Cdr2 or altered Wee1 regulation, we examined a *wee1-2x* mutant carrying an additional copy of *wee1* at an exogenous locus (Figure 4E). Interestingly, *wee1-2x* cells increased their SA:Vol ratio under all stress conditions (Figure 4F). These cells decreased in length under oxidative, osmotic, and low glucose conditions, with no change in width except a decrease under osmotic stress (Figure 4G and 4H). Elevated *wee1* levels lead to more inactive Cdk1, causing *wee1-2x* cells to divide at a larger size (Miller *et al*., 2023) (Figure S1A). This enhanced Cdk1 inhibition likely requires additional Cdr2-mediated suppression to trigger mitotic entry and the resulting delay in division may allow for greater surface area accumulation before mitosis.

Together, these findings identify Cdr2 as a key regulator of surface area-based geometric adaptation under glucose limitation. Loss of *cdr2* shifts cells toward volume-based scaling in low glucose, while increased *wee1* enhances surface area accumulation under all conditions. Thus, Cdr2 and Wee1 operate as a surface area-sensing module that complements the volume-sensing (Cdc25) and time-sensing (Cdc13) pathways to maintain geometric homeostasis under environmental stress.

Despite 48 hours of stress exposure, none of the *cdc25, cdc13*, or *cdr2/wee1* mutants showed major defects in growth or viability, as indicated by similar septation index values across conditions (Figure S2). Thus, even when geometric scaling was altered, cell cycle progression remained robust. These results suggest that surface area- or volume-based adaptation is not essential for survival but instead may fine tune growth and recovery during prolonged stress. Future studies tracking cell populations through chronic or recurring stress will clarify how these adaptive geometries shape long-term proliferation, recovery, and evolutionary resilience.

The stress-specific size phenotypes observed here highlight how size control pathways intersect with stress signaling networks. To our knowledge, stress-induced cell size changes have not been systematically characterized for the *cdc25, cdr*2, and *cdc13* mutants examined. We observed that *cdc25* mutants elongated under osmotic and oxidative stress, consistent with Cdc25’s previously reported role in the Sty1 MAP kinase pathway (López-Avilés *et al*., 2005, 2008), which mediates cellular response to these stresses (Millar *et al*., 1995; Shiozaki and Russell, 1995; Degols *et al*., 1996; Shiozaki *et al*., 1998;). *cdr2Δ* cells failed to reduce their size and maximize their surface area under low glucose conditions, suggesting that Cdr2 links glucose availability to mitotic entry. Cdr2 forms cortical nodes whose activity and localization are modulated by the Pom1 gradient, which itself is responsive to glucose availability (Kelkar and Martin, 2015; Allard *et al*., 2019). Although *cdr2Δ* cells were able to maximize their volume under osmotic stress, they did not reduce their length like WT cells, suggesting that Cdr2 may contribute to osmotic stress-dependent division control, potentially through integration of Sty1 signaling. Notably, Sty1 directly regulates the Cdr2-related kinase Cdr1, preventing its colocalization with Wee1 during osmotic stress (Opalko and Moseley, 2017). Consistent with our finding that Cdc13 contributes to size adaptation under osmotic stress, *cdc13* expression is known to be modulated by the Atf1 transcription factor, a downstream effector of Sty1 (Wilkinson *et al*., 1996; Bandyopadhyay *et al*., 2014). The requirement for Cdc13 in osmotic but not oxidative stress suggests stress-specific modulation of Sty1-Atf regulation. Future studies will be needed to determine how stress signaling networks influence scaling of Cdc25, Cdr2, and Cdc13 with volume, surface area, or time, and to distinguish transient stress responses from long-term adaptation.

Collectively, our results show that fission yeast alter their cell geometry by maximizing either surface area or volume depending on environmental conditions. Volume-based adaptation in oxidative stress requires Cdc25, osmotic stress adaptation requires both Cdc25 and Cdc13, and surface area-based adaptation in low glucose relies on Cdr2 (Figure 5). When one pathway is disrupted, cells often switch to an alternate geometric mode, for example, from volume-to surface area-based control, revealing adaptive flexibility within the network. This demonstrates that having an integrated size control system enables cells to dynamically reconfigure how size is sensed and regulated under stress. Future studies should explore how geometric control principles are conserved or differ across cell types and environmental conditions. Defining how conserved stress-responsive pathways (Sty1-Srk1, PKA, TOR) regulate Cdc25-, Cdc13-, and Cdr2-scaling properties will provide insight into how eukaryotic cells tune Cdk1 activity and maintain robust size homeostasis in changing environments.

**Figure 5:**
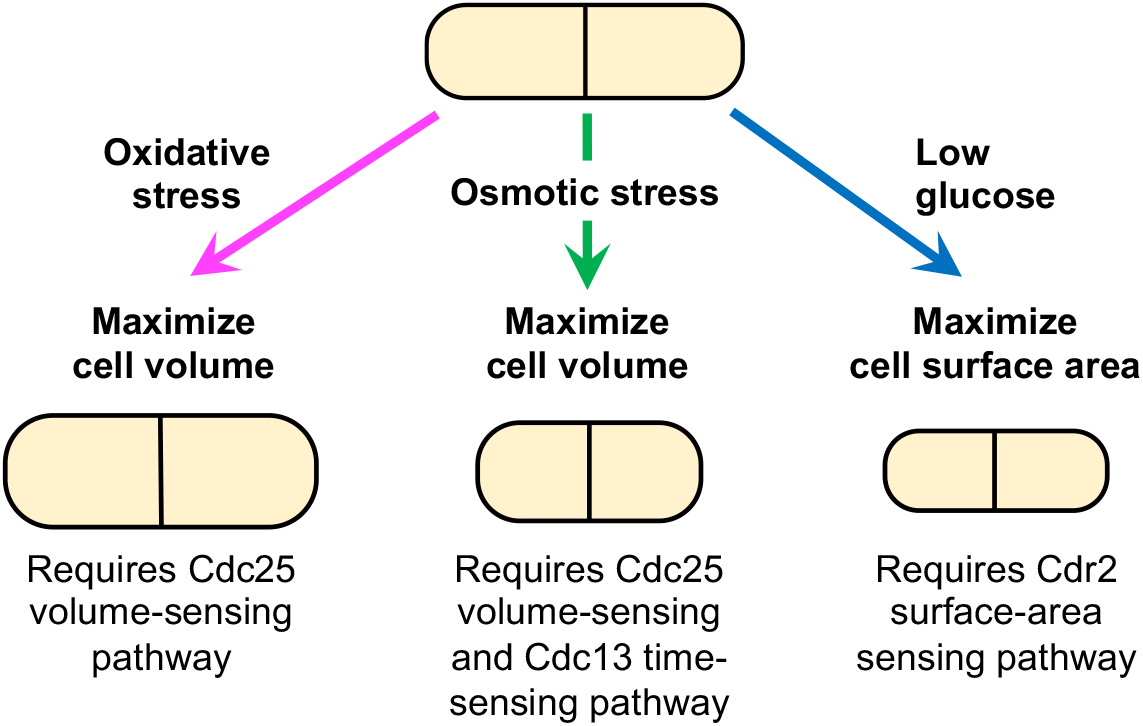
Distinct size control pathways mediate stress-specific geometric adaptations in fission yeast. Cdc25, Cdc13, and Cdr2 mediate distinct geometric responses to oxidative, osmotic, and low glucose stress, enabling fission yeast to flexibly adapt cell size for robust division under changing environments.

## MATERIALS AND METHODS

### Strain construction and growth conditions

Standard methods were used to grow *Schizosaccharomyces pombe* cells (Moreno *et al*., 1991). Yeast strains used in this study are listed in Supplemental Table S1. Gene fusions were expressed under their endogenous promoters unless noted. PCR-based homologous recombination was performed for tagging of genes on the chromosome (Bähler *et al*., 1998). To place *cdc25* under the control of the constitutive *eno101* promoter, we synthesized a gBlock fragment (Integrated DNA Technologies) containing the *kanMX6* marker and a 276 bp *eno101p* sequence (Wang *et al*., 2014). The *kanMX6-eno101p* cassette was PCR-amplified and integrated at the 5′ end of the *cdc25* open reading frame on the chromosome using standard N-terminal tagging methods (Bähler *et al*., 1998). This strain was confirmed to have significantly reduced size-dependent accumulation of Cdc25 by analysis of fluorescent protein intensity, described below. To obtain *cdc13-2x* cells carrying an additional copy of *cdc13+*, we integrated the *StuI*-linearized *pHis5StuI-cdc13-sfGFPint* plasmid (a kind gift of Jamie Moseley) at the *his5* locus of *his5-D21* yeast cells. This strain was confirmed to be correct by growth on EMM -his plates and detection of Cdc13-sfGFPint signal by fluorescent microscopy.

Cells were cultured in standard YE4S medium (3% glucose) overnight and then maintained or shifted to stress conditions. Oxidative stress was induced by supplementing YE4S with 1 mM hydrogen peroxide, osmotic stress with 1 M KCl, and glucose limitation with 0.08% glucose. Cultures were incubated at 25 °C while shaking at 180 rpm for 48 h prior to imaging to allow cells to acclimate to stress conditions. Each yeast strain was grown in triplicate under each condition to ensure reproducibility of cell size measurements. After confirming consistency across individual trials, data sets were combined and plotted together.

### Microscopy

#### Analysis of cell geometry from static images

Cells were harvested by brief centrifugation, resuspended, and ∼2 µl of concentrated suspension was spotted onto 35-mm glass-bottom dishes (P35G-1.5-20C; MatTek) overlaid with YE4S agar (prepared with the same condition as the culture medium and prewarmed to 25°C). Imaging was performed using a widefield epifluorescence microscope: Nikon Ti2-A inverted microscope (NIS-Elements software) equipped with a Prior Z-motorized focus drive; a 60× 1.4 NA CFI60 Plan Apochromat Lambda D oil-immersion objective lens (Nikon); a D-LEDi fluorescence LED illumination system with DAPI, GFP, and mCherry filter cubes; and a PCO.panda USB 3.1 sCMOS camera. For each cell type and condition, multiple fields of view were imaged. Brightfield and DAPI images were captured with 27 optical z-sections and 0.2-µm step size. Cell images displayed in figures are brightfield single z-section images and scale bars are 5 μm.

#### Analysis of fluorescent protein intensity from static images

For imaging to examine size-dependent accumulation of *eno101p-cdc25-mNeonGreen* compared to WT *cdc25-mNeonGreen*, cells were mounted on a coverglass-bottom dish (P35G-1.5-20C; MatTek) and covered with a piece of YE4S agar prewarmed to 25°C. Imaging was performed using a spinning disk confocal microscope (Yokogawa CSU-WI; Nikon Software) equipped with a 60× 1.4-NA CFI60 Plan Apochromat Lambda D oil-immersion objective lens (Nikon); 405-, 488-, and 561-nm laser lines; and a Photometrics Prime BSI camera on an Eclipse Ti2 inverted microscope (Nikon). Multiple fields of view were acquired for each cell type, with 27 optical z-sections collected at 0.2-µm intervals.

### Cell and nuclei segmentation

Cell and nuclei segmentation was carried out using a semi-automated pipeline as described in Miller *et al*., 2023. In brief, nuclear segmentation was based on the BFP-NLS signal, which reliably marked nuclei while Brightfield images were used for cell segmentation. In ImageJ, Brightfield stacks were smoothed with Gaussian filtering, followed by global thresholding to generate binary masks of cell outlines. Morphological operations (erosion and dilation) were used to refine masks and separate adjacent cells. Artifacts, edge cells, and unresolved clumps of cells were removed manually in ImageJ. The resulting binary image (“cell mask”) was compared with the original brightfield image and confirmed to be an accurate representation of cell size.

BFP-NLS images were processed for nuclear segmentation using a semi-automated ImageJ pipeline (as in Miller *et al*., 2023). In ImageJ, a sum projection was created from z-stacks and were subsequently smoothed, and globally thresholded to generate binary nuclear masks. Masks were filtered to exclude edge objects. To correct for uneven BFP illumination across an imaging field, abnormally small or large nuclei were removed from binary masks. The resulting binary masks (‘nuclei mask”) was then validated against the original BFP-NLS images to ensure accurate nuclear size representation.

Cell masks of dividing cells were generated using ImageJ and MATLAB as described in Miller *et al*., 2023. Briefly, nuclear masks were eroded in ImageJ to separate nuclei from cell borders, then combined with cell masks in MATLAB to produce nuclei-overlaid images. These were filtered to retain only cells containing two nuclei, manually confirmed, and further processed to remove nuclei, yielding binary masks of dividing cells.

### Cell geometry measurements

Cell geometry measurements were obtained as described in Miller *et al*., 2023. In brief, cell widths (≥50 cells per condition) were manually measured from cell masks in ImageJ using the straight-line tool, avoiding irregular regions. This approach provided more reproducible measurements for surface area and volume calculations than automated methods. For each strain and condition, the mean cell radius (average width/2) was used to calculate cell surface area and volume, assuming a uniform radius across the population (Table 1). Cell length or symmetry axes were identified in MATLAB by applying principal component analysis of the cloud points within each segmented cell (Facchetti *et al*., 2019). Cell surface area and volume were calculated in MATLAB using the formulas for a cylinder with hemispherical ends, reflecting the rod-like geometry of fission yeast (Table 1). Cell volume and surface area values from our experiments were consistent with previously published work (Facchetti *et al*., 2019; Miller *et al*., 2023).

**Table 1.**
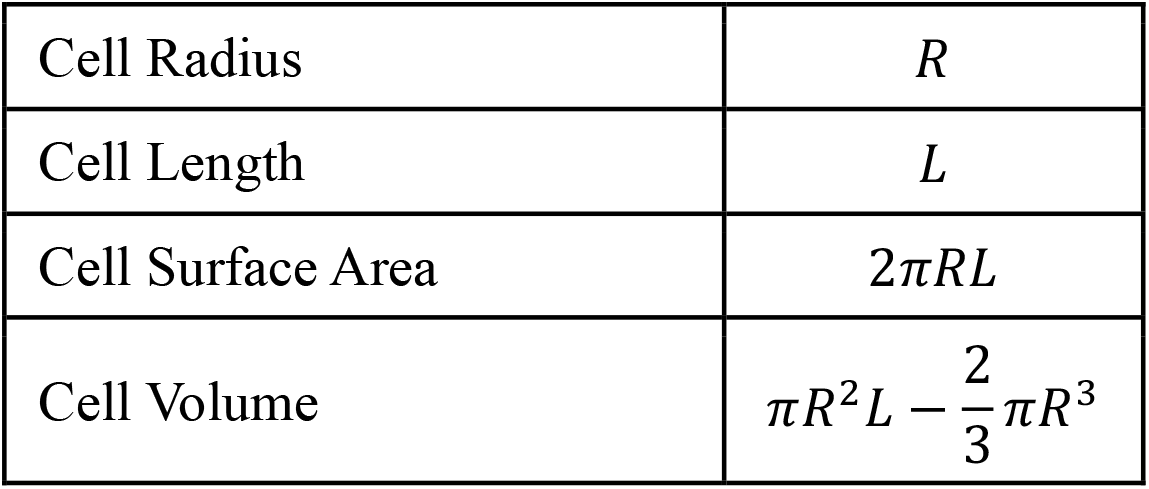
Calculation of cell surface area or volume.

|  |  |
| --- | --- |
| Cell Radius | $R$ |
| Cell Length | $L$ |
| Cell Surface Area | $2\pi RL$ |
| Cell Volume | $\pi R^2 L - \frac{2}{3}\pi R^3$ |

### Measurement of fluorescent intensity

Summed projections of z-sections were used to analyze fluorescent intensity. MATLAB scripts quantified mean fluorescence of proteins in the cytoplasm, nucleus, or whole cell, as defined by cell and nuclear masks.

All MATLAB codes used for image processing and analysis are available on GitHub (https://github.com/millerk89/sizer-timer-pombe), as previously archived in Miller *et al*., 2023.

### Septation index analysis

Brightfield images of cells, taken from the same data sets used for cell size measurements, were analyzed to determine septation index. For each strain and condition, at least 100 cells were counted to calculate the percentage of cells with a single septum vs. no septum.

### Statistical analysis

Data analysis was performed in GraphPad Prism. Data are presented as mean values with standard deviation unless otherwise noted. Comparisons among multiple strains or growth conditions were assessed by ANOVA. Comparisons between two groups were assessed using an unpaired two-tailed t-test. p-values < 0.05 were considered statistically significant. Linear regression analysis was used to obtain slope values for assessing relationship between cell volume and Cdc25 mean intensity in the nucleus for WT and *eno101p-cdc25* cells.

## Abbreviations

WT: wild-type
SA: surface area
Vol: Volume
SA:Vol: SA-to-Vol
mNG: mNeonGreen
NLS: Nuclear localization sequence
BFP: blue fluorescent protein.

## ACKNOWLEDGEMENTS

We thank members of the Miller lab for helpful discussions or comments on the manuscript; and James B. Moseley for sharing yeast strains and plasmids. This work was funded by grants from the NIH RI-INBRE (2P20GM103430, ECD subaward 0012496/0603246) and the Rhode Island Foundation (#16411_139175) to K.E.M. This research was also supported by a NIH S10 Instrumentation Grant (1S10OD038193-01) to K.E.M, which funded the Nikon CSU-W1 Spinning Disk Confocal Microscope. We also acknowledge research support through a Walsh Fellowship from Providence College awarded to E.C.

## FIGURE LEGENDS

**Table S1.** Yeast strains used in this study.

| Strain | Genotype | Source |
| --- | --- | --- |
| KMY127 | *lys3+::ptdh1\*:NLS-linker-mTagBFP2:terminatordh1:nat R ULA+ h-* | Lab collection |
| KMY357 | *cdr2Δ::NAT lys3+::ptdh1\*:NLS-linker-mTagBFP2:terminatordh1:kanMX ULA+* | Lab collection |
| KMY378 | *(AKA wee1-2x) lys3+::ptdh1\*:NLS-linker-mTagBFP2:terminatordh1:kanMX leu1-32:[pJK148-Pwee1-wee1-Twee1] UA+* | (Miller *et al.*, 2023) |
| KMY368 | *kanMX6::eno101p-cdc25-mNeonGreen::hphR lys3+::ptdh1\*:NLS-linker-mTagBFP2:terminatordh1:natR h- ULA+* | This study |
| KMY376 | *cdc25-degron-DAmP::kanMX6 lys3+::ptdh1\*:NLS-linker-mTagBFP2:terminatordh1:natR ULA+ h* | Lab collection |
| KMY385 | *(AKA cdc13-2x) pHis5StuI-cdc13-sfGFPint lys3+::ptdh1\*:NLS-linker-mTagBFP2:terminatordh1:natR ULA+* | This study |
| KMY349 | *his5-D21 h-* | Vještica *et al.*, 2020 |

**Figure S1:**
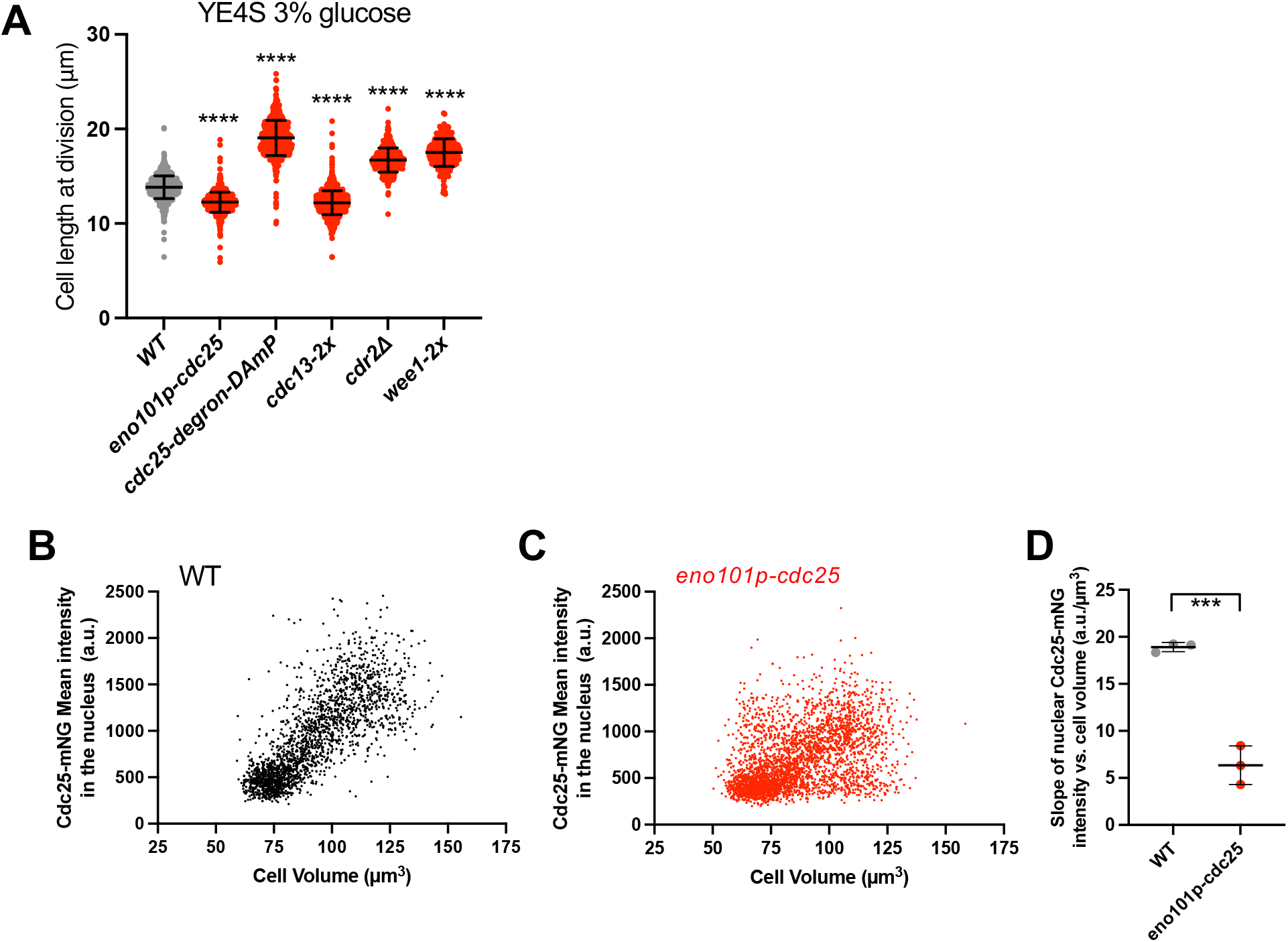
Verification of fission yeast strains by cell length and fluorescence. (A) Cell length at division of indicated strains grown in 3% glucose YE4S media. All \*\*\*\**p* < 0.0001 compared to WT. (B) Cdc25-mNG nuclear mean intensity in WT cells; n=2,133 or (C) *eno101p-cdc25* cells; n=3,160. (D) Slopes from linear regression of three independent trials examining Cdc25 nuclear mean intensity vs. cell volume in WT or *eno101p-cdc25* cells. Each trial n> 500 cells. \*\*\**p* =0.0005.

**Figure S2:**
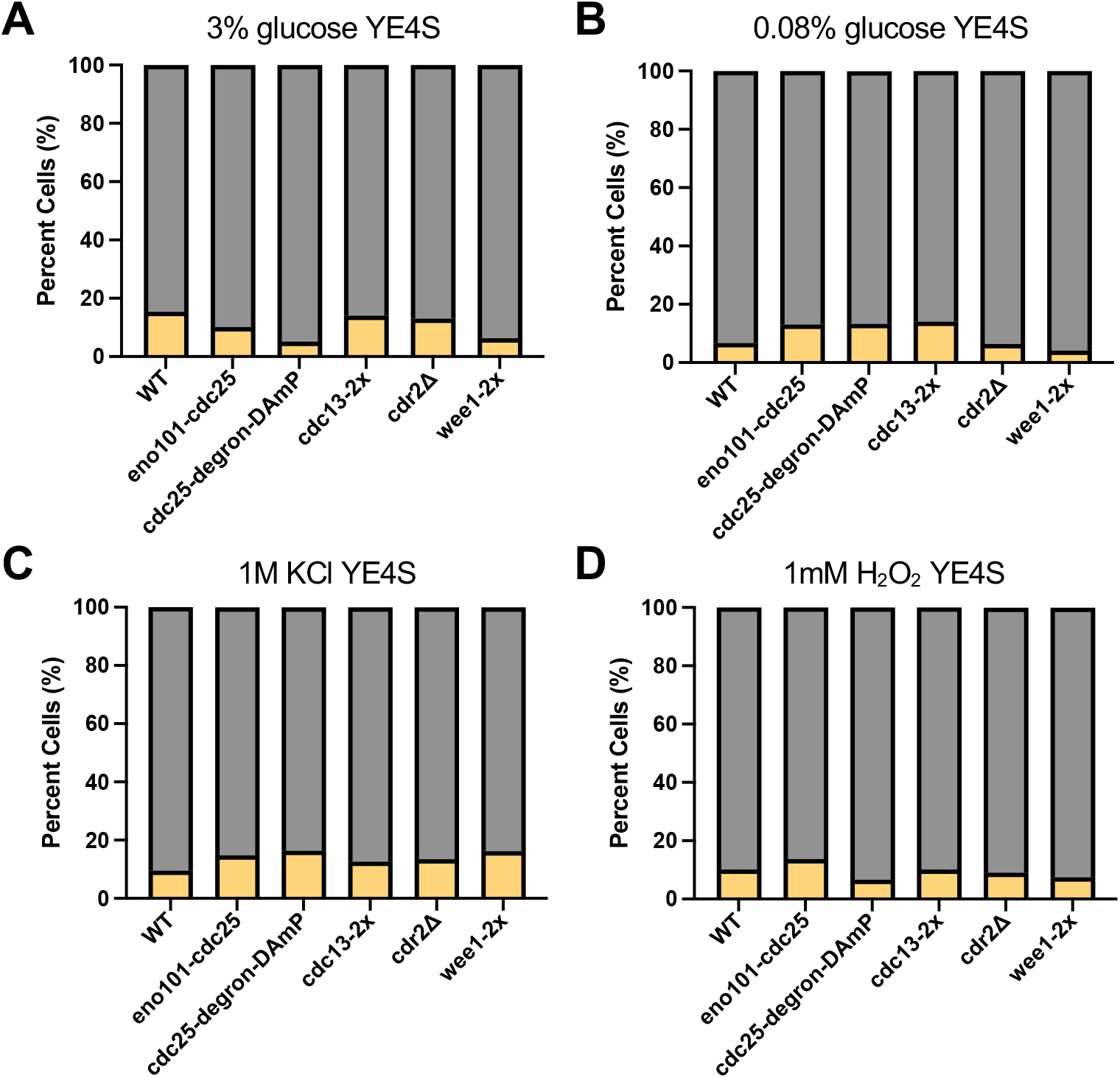
Septation index of fission yeast strains grown under various conditions. (A) Yellow bar, single septum; Grey bar, no septum. n ≥ 100 cells per strain and condition.

